# How does the ion concentration affect the functions of kinesin BimC

**DOI:** 10.1101/2024.05.31.596855

**Authors:** Wenhan Guo, Yuan Gao, Dan Du, Jason E Sanchez, Akasit Visootsat, Yupeng Li, Weihong Qiu, Lin Li

## Abstract

BimC family proteins are bipolar motor proteins belonging to the kinesin superfamily which promote mitosis by crosslinking and sliding apart antiparallel microtubules. Understanding the binding mechanism between the kinesin and the microtubule is crucial for researchers to make advances in the treatment of cancer and other malignancies. Experimental research has shown that the ion concentration affects the function of BimC significantly. But the insights of the ion-dependent function of BimC remain unclear. By combining molecular dynamics (MD) simulations with a series of computational approaches, we studied the electrostatic interactions at the binding interfaces of BimC and the microtubule under different KCl concentrations. We found the electrostatic interaction between BimC and microtubule is stronger at 0 mM KCl compared to 150 mM KCl, which is consistent with experimental conclusions. Furthermore, important salt bridges and residues at the binding interfaces of the complex were identified, which illustrates the details of the BimC-microtubule interactions. Molecular dynamics analyses of salt bridges identified that the important residues on the binding interface of BimC are positively charged, while those residues on the binding interface of the tubulin heterodimer are negatively charged. The finding in this work reveals some important mechanisms of kinesin-microtubule binding, which helps the future drug design for cancer therapy.

## Introduction

The BimC family proteins belong to the kinesins’ superfamily, which plays vital roles in transporting various cargos for different cellular processes [1]. BimC family proteins are bipolar motor proteins that exert their functions by crosslinking and sliding apart antiparallel microtubules [2]. Bipolar BimC proteins include those found in *Aspergillus nidulans* (BimC), *Saccharomyces cerevisiae* (CIN8), *Schizosaccharomyces pombe* (CUT7), *Drosophila* (KLP61F), *Xenopus* Eg5 (XlEg5), and humans (HsEg5) [3–9]. The subfamily members have similar N-terminal motor domains and contain a conserved BimC box region in the C-terminal tail domain [6, 10–12].

Many experimental studies have been carried out to determine the function and motility of BimC family proteins [8, 13–17]. In particular, mutating residues on BimC is an important way to study the function of BimC and helps to understand other kinesins, including Eg5, an anti-cancer drug target [5, 11]. Dr. I.M. Hagan’s group mutated residues on the Cut7 BimC box region and found divergence of regulation within the BimC family [11]. Another work performed by TJ Mitchison demonstrated that a single point mutation in the most conserved part of the BimC box is capable of disrupting localization to the mitotic spindle [18]. Dr. N. Ronald Morris’s group found that a mutation of the gene that encodes the BimC protein blocks nuclear division in *Aspergillus nidulans* [5].

Computational methods also have been widely used to study BimC proteins. Dr. Endow’s group at Duke University Medical Center built the kinesin phylogenetic tree by parsimony methods from sequence alignment approaches [19]. Dr. Sims’s group studied the inhibition of the HsEg5 and other homologues in the BimC subfamily using molecular docking and molecular dynamics approaches, which could potentially aid in drug design [20]. Dr. Francois Delfaud and Dr. Fabrice Moriaud’s groups explored the hydrophobic sub-pocket of the mitotic kinesin Eg5 allosteric binding site using computational fragment-based drug design through the MED-SuMo software [21].

Previous work in our laboratory has shown that electrostatic features on the motor domains of Eg5 provide attractive interactions with the microtubule. Additionally, on the binding interfaces of Eg5 and the tubulin heterodimer, salt bridges play the most significant role in holding the complex, which indicates an asymmetric binding mechanism whereby Eg5 moves along the microtubules [9].

In this work, we focus on exploring BimC motility on microtubules. By using experimental approaches and muti-scale computational methods including DelPhi [22], DelPhiForce [23, 24], and NAMD [25], the binding mechanism (specifically, key electrostatic features) of BimC and the microtubule was investigated. Single molecule experiments were conducted to compare the interactions between BimC and microtubules at different ion concentrations, which demonstrated the motility features of BimC are quite different when the KCl concentration varies from 0 to 150 mM. Computational studies of BimC under 0, and 150 mM KCl concentrations were studied to reveal the mechanisms of BimC motility features at different ion concentrations.

The electrostatic forces between the BimC motor domain and tubulin heterodimer under 0, and 150 mM KCl ion concentrations were calculated. The results of calculated electrostatic forces were consistent with experiment results: higher ion concentration results in lower binding affinity. The 150 mM KCl concentration yields a very low binding affinity, which is consistent with our experimental results: At such ion concentration the BimC doesn’t even bind to microtubules in the experiment. Then the electrostatic potential on the binding surfaces of the BimC motor domain and tubulin heterodimer were shown to further support this finding. Finally, important salt bridges were identified with MD simulations. Several of these residues are identified to be critical for BimC binding to the microtubule, and characterization of these can help guide kinesin-focused anticancer drug design [26–29].

## Methods

### Generating recombinant BimC(Δ1-70)-GFP construct

cDNA of full length BimC was codon optimized for protein expression in *E coli*. and then synthesized by IDT. Recombinant BimC plasmid was generated by integration of the BimC cDNA fragment into a modified Novagen pET-17b vector containing a 6xHis-tag and a GFP using Gibson Assembly (NEB). Plasmid region corresponds to the 71-1184 amino acids of BimC and the modified vector was amplified and assembled with KLD Enzyme Mix Reaction (NEB) to generate the recombinant BimC(Δ1-70)-GFP construct. Finally, the recombinant construct was verified by DNA sequencing (GENEWIZ).

### BimC(Δ1-70)-GFP protein expression and purification

Recombinant plasmid containing BimC(Δ1-70)-GFP was transformed to and expressed in Novagen Rosetta (DE3) competent cells. Cells were grown at 37 °C in tryptone phosphate medium (TPM) supplemented with 50 μg mL-1 ampicillin. Protein expression was induced by 0.1 mM IPTG on ice when OD600 reached 0.8. After incubation for an additional 12∼14 h at 18 °C, cell pellet was harvested by centrifugation at 4,550 g for 30 min using a JS-4.2 (Beckman Coulter). For protein purification, cell pellets were resuspended in the lysis buffer (50 mM sodium phosphate buffer (pH 7.2) with 500 mM NaCl, 1 mM MgCl2, 0.5 mM ATP, 10 mM β-mercaptoethanol, 20 mM imidazole, and protease inhibitor cocktail), lysed via sonication and centrifuged at 27,200 g for 30 min using a JA-20 (Beckman Coulter). Soluble protein in the supernatant was purified by Talon metal affinity resin (Takara Bio) and eluted into the elution buffer (50 mM sodium phosphate buffer (pH 7.2) with 500 mM NaCl, 1 mM MgCl2, 0.5 mM ATP, 10 mM β-mercaptoethanol, 250 mM imidazole). Finally, protein was flash-frozen in liquid nitrogen and stored at −80 °C.

### Preparation of polarity-marked HiLyte 647 microtubules

Taxol-stabilized polarity-marked HiLyte 647 microtubules with bright plus-ends were prepared as previously described [30]. Firstly, a dim tubulin mix containing 17 mM unlabeled tubulin, 0.8 mM HiLyte 647 tubulin and 17 mM biotinylated tubulin was first incubated in BRB80 with 0.5 mM GMPCPP at 37 °C overnight (16h) for the dim microtubule polymerization, and then centrifuged at 250,000 g for 7 min at 37 °C in a TLA100 rotor (Beckman). The pellet was then resuspended in a bright tubulin mix containing 7.5 mM unlabeled tubulin, 4µM HiLyte 647-tubulin, and 15µM NEM-tubulin in BRB80 with 2 mM GMPCPP and incubated at 37 °C for 50 min to cap the plus-end of the dim microtubules. Finally, polarity-marked microtubules were pelleted at 20,000 g for 7 min at 37 °C in the TLA100 rotor (Beckman Coulter) and resuspended in BRB80 buffer with 40 mM taxol.

### Total internal reflection fluorescence (TIRF) microscopy

All TIRF microscopy experiments were performed at room temperature using the Axio Observer Z1 objective-type TIRF microscope (Zeiss) equipped with a 100x 1.46 NA oil-immersion objective and a back-thinned electron multiplier CCD camera (Photometrics). Flow chambers were made by attaching a coverslip to a microscope glass slide by a double-sided tape. All experiments used coverslips functionalized with Biotin-PEG to reduce the nonspecific surface absorption of protein molecules as previously described [31].

### In Vitro Single-Molecule Motility Assay

In single molecule assays, flow chamber was perfused with 0.5 mg/mL streptavidin for immobilizing polarity-marked HyLite 647 microtubules. Purified BimC(Δ1-70)-GFP protein was diluted in the motility buffer (BRB50 supplemented with 1 mM ATP, 10 mM KCl, 25 mM taxol, 1.3 mg mL-1 casein, and an oxygen scavenger system [32]. Time-lapse images were acquired at 1 frame per second with 100-ms exposure time and 5-min recording time. Kymographs were generated and analyzed in ImageJ (NIH) for determining directionality, velocities, and run-length of BimC(Δ1-70)-GFP molecules. Velocity and run length were determined by fitting the histograms to a Gaussian distribution and an exponential distribution respectively using SciPy.

### Computational structure preparation

The experiments of our work showed that the binding affinity of BimC and the microtubule decreases when the ion concentration is increased. The experimental work shows there’s no bind between BimC and microtubules at 150 mM KCl concertation. While BimC(Δ1-70)-GFP at 0mM KCl condition shows active behavior. BimC walking to the minus end along the microtubule with high velocity. To explore the detailed binding mechanism of BimC and the microtubule, we modeled the motor domain of BimC (residues 71 to 413) by AlphaFold2 [33, 34]. In the BimC structure, the N-terminal domain (with residues 1 to 70) is intrinsically disordered; therefore, the N-terminal domain of BimC was not included in this work. Finally, the complex structure of BimC binding to a tubulin heterodimer was modeled based on PDB entry 6TA4 [35], which is the motor domain of Eg5 in the adenylyl imidodiphosphate (AMP-PNP) state bound to tubulin proteins at a 6.10 Å resolution.

### MD Simulations

The complexes described above were explored with NAMD[25] using an explicit solvent model. MD simulations was carried out on Stampede2 at the Texas Advanced Computing Center (http://www.tacc.utexas.edu). A 20 ns simulation was performed on the BimC-tubulin heterodimer complexes at 0/150 mM KCl (see Supplementary Movies 3-4). In the simulations, the minimization was set to 20,000 steps, the temperature was set to 300 K, and the pressure model is set as Langevin dynamics. The full-system periodic electrostatics coordinates fit the grid size (100, 70, 100). Residues with any atom within 10 Å from the binding interfaces were treated as interfacial residues. All the interfacial residues were set free while non-interfacial residues were constrained. Based on the Root Mean Square Deviation (RMSD) plot (see Supplementary Figure S1), 2000 frames from the last 10 ns (from 10 ns to 20 ns with 2000 frames) of the simulations were selected for analysis. The simulations were visualized by VMD [36] (see Supplementary Movies 3-4). To further explore the interactions between the BimC motor domain and the tubulin heterodimer, the salt bridges between the two interfaces at 0 mM KCl concentration were identified and analyzed using the salt bridge extension in VMD[36]. The threshold for salt bridges was set to 4.0 Å. The salt bridges (with occupancy >30%) and the important interfacial residues (with occupancy >66%) that contribute significantly to the binding interactions during the MD simulation, was identified and explored.

### Electrostatic Force Calculations

Experimental results show that BimC processively walks to the minus end on microtubule at 0 ion concertation, while BimC doesn’t bind to microtubules at 150 mM KCl concertation. To compare the results of our calculations to the conclusions from the experiment and investigated the mechanism between BimC motor domain and microtubule, DelPhiForce [23, 24] was used to calculate the electrostatic binding forces between the BimC and tubulin heterodimer. The tool calculates electrostatic forces by solving the Poisson-Boltzmann equation (PBE) using the finite difference method:

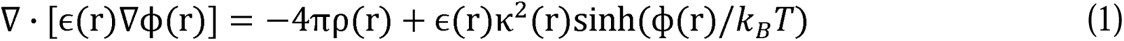

where ϕ(r) is the electrostatic potential, ρ(r) is the permanent charge density, ε(r) is the dielectric permittivity, *κ* is the Debye-Hückel parameter, *k_B_* is the Boltzmann constant, and *T* is temperature.

The complex structures used to calculate the forces were extracted from the two different simulations. To be precise, we extracted the structure of the 4000th frame (the last frame in the simulations) from the 0 and 150 mM KCl simulations. The structure in the last frame was chosen for the force calculations because the structure of the complex is reliable after the MD simulation. To study the directions and strengths of the net forces, DelPhiForce was utilized to calculate the electrostatic forces between the BimC motor domain and the tubulin heterodimer at variable distances. The BimC motor domain and the tubulin heterodimer were separated from 10 Å to 40 Å with a step size of 2 Å using StructureMan [37]. The net forces were displayed using Visual Molecular Dynamics (VMD) [36] as shown in Figures 1-2.

**Figure 1:**
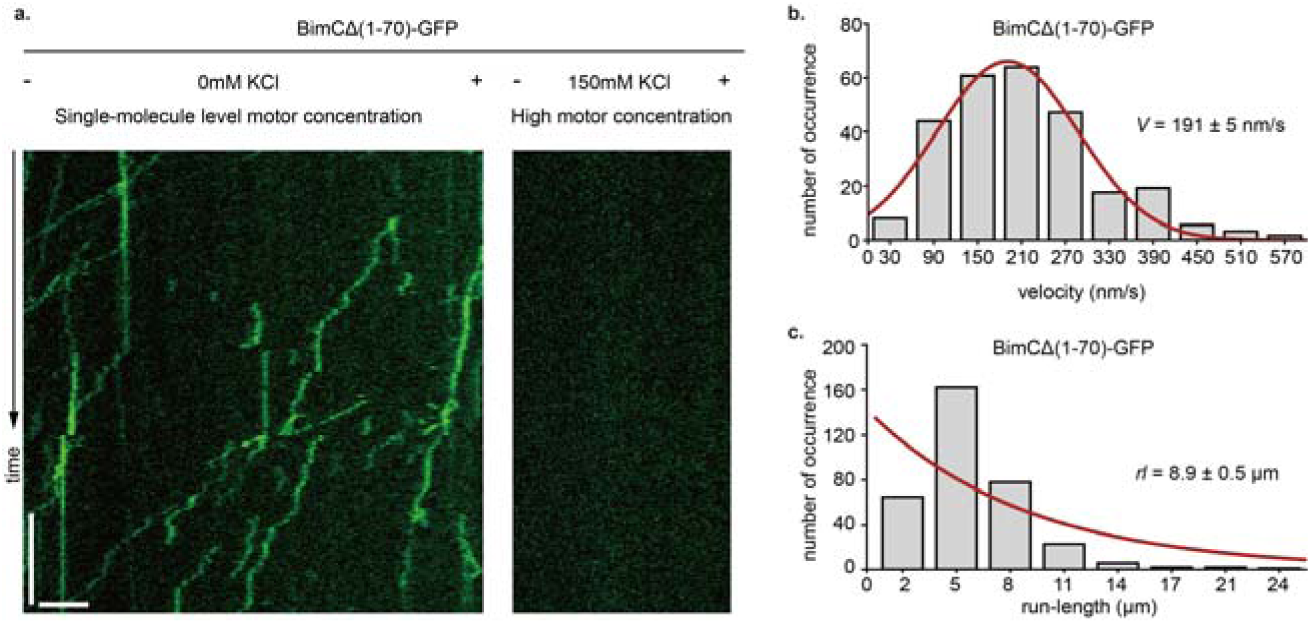
BimC(Δ1-70)-GFP construct’s behavior at 0/150 mM KCl concentration. (a) Left panel: Representative kymograph of BimC(Δ1-70)-GFP at 0mM KCl condition and single-molecule level motor concentration. Right panel: Representative kymograph of BimC(Δ1-70)-GFP at 150mM KCl condition and very high motor concentration. Scale bars: 1 minute (vertical) and 5 µm (horizontal). (b) The velocity histogram of BimC(Δ1-70)-GFP at 0mM KCl condition (n = 339). The red curve represents a Gaussian fit. (c) The run length histogram of BimC BimC(Δ1-70)-GFP at 0mM KCl condition (n = 339). The red curve represents an exponential fit.

**Figure 2.**
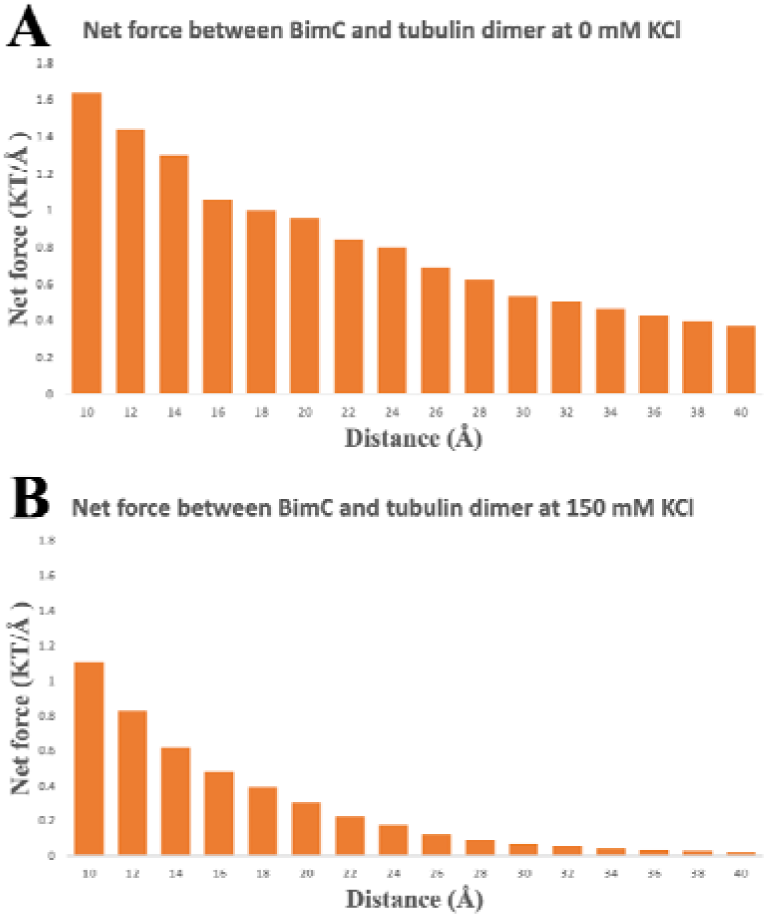
The electrostatic forces between the BimC motor domain and tubulin heterodimer at distances from 10 Å to 40 Å with a step size of 2 Å at 0/150 mM KCl concentrations.

### Electrostatic potential Calculations

To further investigate the interaction of the BimC motor domain and tubulin heterodimer, DelPhi [22] was utilized to calculate the electrostatic potential of the complex under 0 and 150 mM KCl concentrations. DelPhi calculates the electrostatic potential of biomolecules by solving the Poisson-Boltzmann equation (PBE) as shown in equation (1). The protein filling percentage of the Delphi calculation box was set to 70.0, the probe radius for the molecular structure was 1.4 Å, and the boundary condition for the Poisson Boltzmann equation was set to the dipolar boundary condition. The calculated electrostatic surface potential was visualized with USCF Chimera [38].

## Results and Discussions

The experimental works were done including generating BimC(Δ1-70)-GFP construct and the representative kymograph of BimC(Δ1-70)-GFP at 0/150 mM KCl condition. To study the mechanisms of the BimC binding with microtubules, we first investigated the electrostatic features of the BimC motor domain and tubulin heterodimer. Furthermore, the salt bridges at different ion concentrations were investigated based on MD simulations. The salt bridges and important residues involved in forming salt bridges between the BimC motor domain and the tubulin heterodimer were analyzed.

### High salt condition negatively impacts the binding affinity between BimC motor domains and microtubules

We set out to determine the behavior of binding affinity between BimC motor domains and microtubules under different salt condition with in vitro single molecule assays. To address the exclusion of the first 70 amino acids in our structural prediction of BimC motor domain, the recombinant BimC(Δ1-70)-GFP construct was generated through molecule cloning with a GFP attached to its C-terminus for visualization. At low-salt condition, BimC(Δ1-70)-GFP bound to microtubules tightly and exhibited ability to walk to the minus end of the microtubule in a processive manner (Figure 1a left panel; Supplementary Movie 1). Statistical analysis revealed an average velocity of 191 ± 5 nm/s (Figure 1b) and an average run length of 8.9 ± 0.5 nm/s (Figure 1c), indicating a steady interaction between the BimC motor domain and microtubules. In contrast, BimC(Δ1-70)-GFP barely landed on the microtubules at high-salt condition, even with a high motor concentration (Figure 1a right panel; Supplementary Movie 2). Collectively, these results suggest that the binding affinity between BimC motor domains and microtubules is negatively impacted by the increase of ionic strength in the environment.

### Electrostatic Forces

DelPhiForce [23, 24] was utilized to study the electrostatic features of the BimC motor domain and tubulin heterodimer. To avoid clashes between BimC and tubulins, the BimC motor domain was separated from the tubulin heterodimer from 10 Å to 40 Å with a step size of 2 Å using StructureMan[39]. Then the electrostatic forces between BimC and the tubulin heterodimer were calculated at each position by DelPhiForce [23, 24].

The magnitudes of the net binding forces between BimC and the tubulin heterodimer at different distances are presented in Figure 2. As shown in Figure 2, for two KCl concentrations, the attractive force decreases when the distance between the motor domain and tubulin heterodimer is increased, which is expected due to Coulomb’s law. Among the different KCl concentrations, the BimC/tubulin heterodimer complex which had the strongest net force was the 0 mM KCl concentration (which is 1.64 KT/Å). The electrostatic force of the BimC/tubulin heterodimer complex was weakest at 150 mM KCl concentration, which is 1.11 KT/Å. This corroborates with our experiment results, which showed a decrease in motor landing rate when the salt condition was increased from 0mM KCl to 150mM KCl (Figure 1a; Supplementary Movie 1; Supplementary Movie 2). In addition, this is consistent with the study of salt adsorption by molecular complexes [40]. In the salt-free state (0 mM KCl concentration), the electrostatic strength between the BimC motor domain and tubulin heterodimer decays with the distance between them. If the concentration exceeds a certain threshold, like 150 mM KCl concentration, the BimC /tubulin dimer complex can be "dissolved". This is due to the shielding of electrostatic interactions between the charged units.

Besides the magnitudes of the electrostatic forces between the BimC motor domain and tubulin heterodimer, the directions of the net forces between the two are also shown for better visualization in the form of arrows. The blue arrows in Figure 3 show the directions of net forces between the BimC motor domain and the tubulin heterodimer at different concentrations. The arrows only represent the directions of the forces because the blue arrows are normalized to the same size for better visualization. Figure 3 reveals that the net forces are attractive between the BimC motor domain and tubulin heterodimer at different concentrations. In the salt-free state (0 mM KCl), the directions of electrostatic forces between the BimC motor domain and tubulin heterodimer change the least compared to 150 mM KCl.

**Figure 3.**
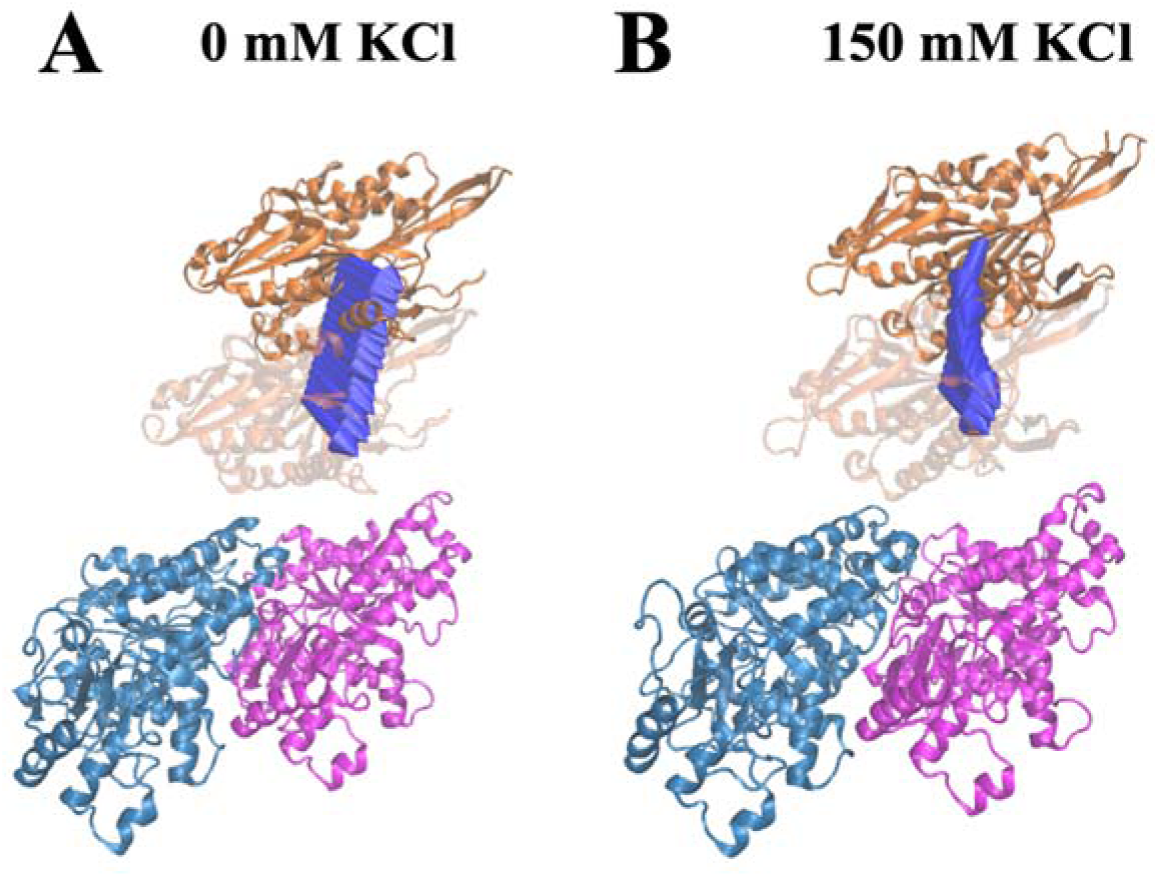
Electrostatic forces of the BimC motor domain and tubulin heterodimer at distances from 10 Å to 40 Å at 0/150 mM KCl concentrations. The α-tubulin, β-tubulin, and BimC units are shown in blue, pink, and orange, respectively. (A) (B) show the directions of the electrostatic forces of the BimC motor domain relative to the tubulin heterodimer at distances from 10 Å to 40 Å with a step size of 2 Å at 0 and 150 mM KCl, respectively.

### Electrostatic potential

The structures and electrostatic potential on the surfaces of the complexes (at 0/150 mM KCl concentrations) are shown in Figure 4. The electrostatic potential is calculated by DelPhi [22] and shown on the surfaces, which are rotated at certain angles to show the electrostatic potential on the binding interfaces. Note here the complex structures used to calculate the electrostatic potential were the structures after MD simulations (at 0/150 mM KCl concentrations). Positively and negatively charged regions are colored in blue and red, respectively. The color scale range is from −3.0 to 3.0 kT/e. The BimC motor domain and tubulin heterodimer are attractive if their interfacial have opposite net charges.

**Figure 4.**
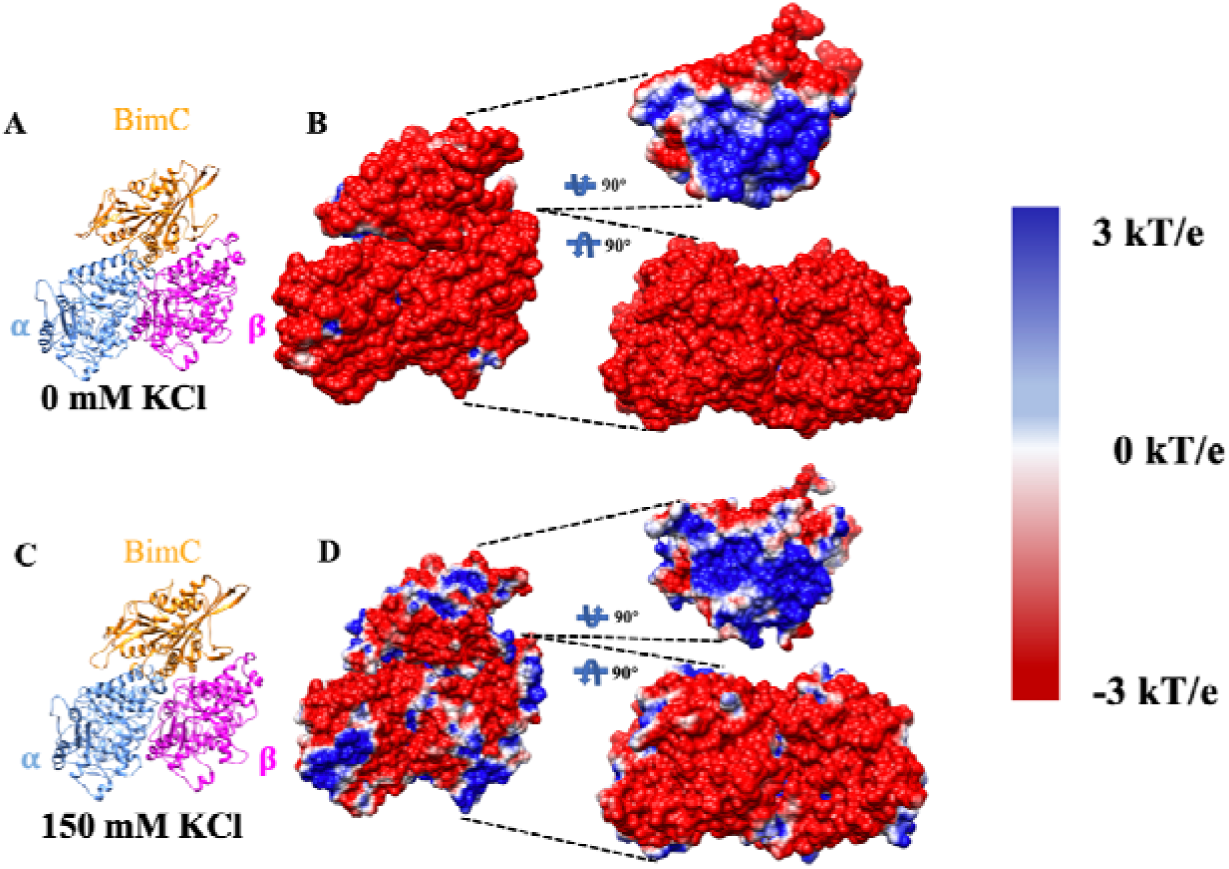
Structures and electrostatic potential of the BimC motor domain and tubulin heterodimer at 0 and 150 mM KCl. (A) (C) are the structures of the complexes after MD simulations at 0 and 150 mM KCl, respectively. (B) (D) are the front views of electrostatic potential for BimC and the tubulin heterodimer which were calculated based on the complex structures after MD simulations. The electrostatic potentials on the binding interfaces were represented in detail by rotating BimC and the tubulin heterodimer 90° in opposite directions.

As shown in Figure 4, at 0 mM KCl concentration, the front view of the complex is dominantly negatively charged. As the concentration of KCl increases, the electrostatic charge in the side view gradually changes from a negative charge dominant to an intertwined positive and negative charge at 150 mM KCl. Compared to the 0 mM KCl structure, the positive and neutral regions in the 150 mM KCl structure increase in size.

Figure 4 also shows the binding interfaces of each structure rotating the BimC motor domains and tubulin protein structures 90° in opposite directions. The BimC motor domain and tubulin heterodimer are attractive if their interfacial have opposite net charges. As shown in Figure 4, the electrostatic potential on the BimC motor domain is primarily positive under the condition of 0 mM KCl. As the KCl concentration increases, the positively and neutrally charged regions on the binding interface of BimC motor domain begin to decrease. As for the tubulin heterodimer, the electrostatic potential on the tubulin heterodimer under 0 mM KCl concentration seems to be prominently negative. As the ion concentration increases, the positively and neutrally charged regions on the tubulin heterodimer binding interfaces begin to increase. In conclusion, the electrostatic potential of the complex under 0 mM KCl has the largest surface area of interacting opposite charges of the two complexes. In the complexes at 150 mM KCl, the binding interfaces of the BimC motor domain and tubulin heterodimer exhibit increased regions of similar charge (either both positively or negatively charged) at the corresponding binding locations, which contributes to the weakening of the electrostatic interaction between the BimC and tubulin heterodimer. Therefore, the BimC motor domain and tubulin heterodimer have the strongest attractive force at 0 mM KCl concentration. The attractive force may enhance the stability of the complex and implies to their tight binding. This finding is consistent with our results from the electrostatic force analysis.

### Salt Bridges

To explore the interaction between the BimC motor domain and tubulin heterodimer in detail, salt bridges with occupancies above 30% on the binding interfaces of the BimC motor domain and α-/β-tubulin at 0 mM KCl are shown in Figure 5.

**Figure 5.**
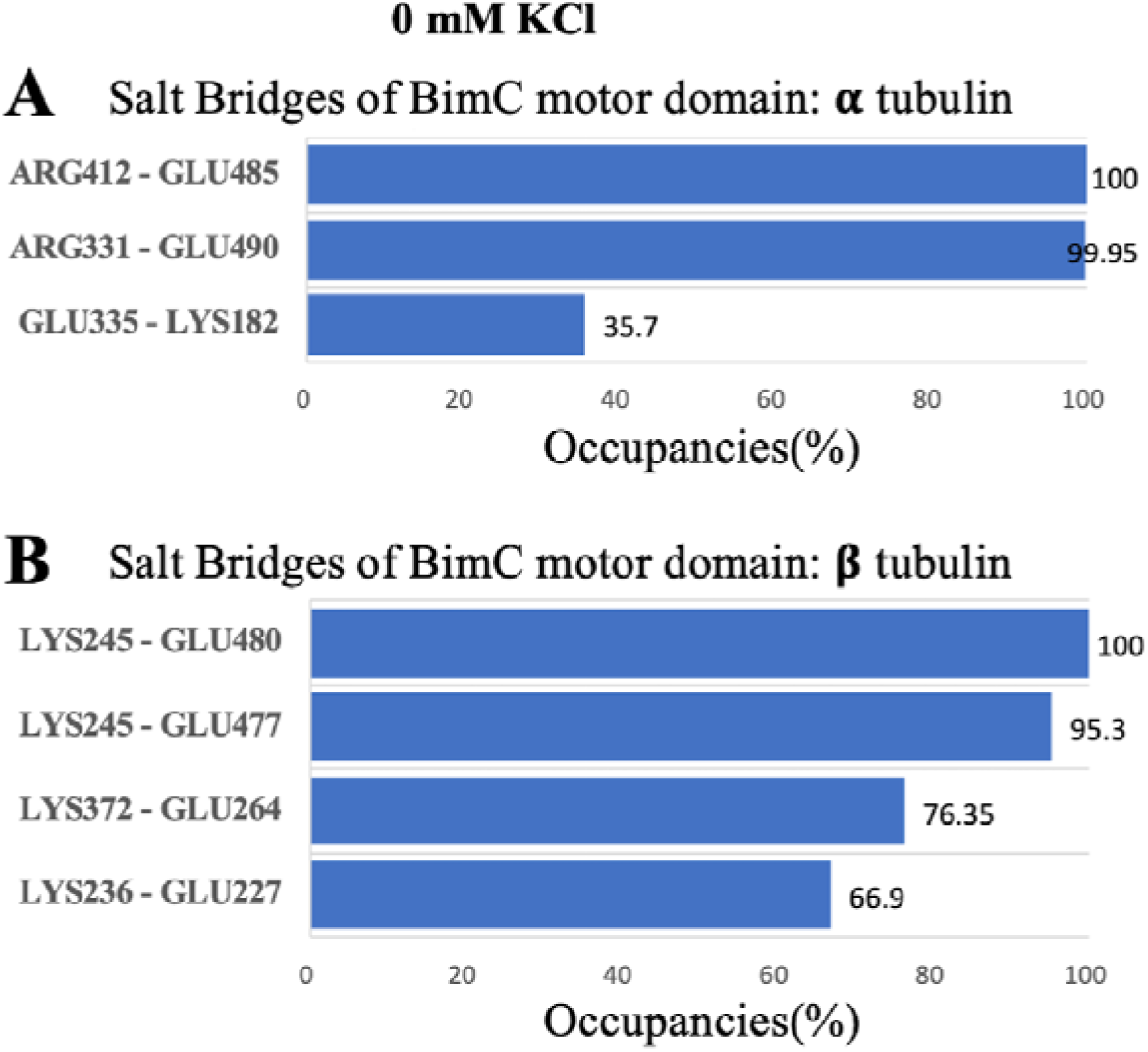
Salt bridges with occupancy above 30% on the binding interfaces of the BimC motor domain and α/β-tubulin at 0 mM KCl.

On the binding interface of BimC and α-tubulin, two pairs of salt bridges, ARG412 (BimC) – GLU485 (α-tubulin) and ARG331 (BimC) – GLU490 (α-tubulin), had occupancies higher than 95% at 0 mM KCl ion concentration. The other two high occupancy salt bridges, LYS245 (BimC) – GLU480 (β-tubulin) and LYS245 (BimC) – GLU477 (β-tubulin), were found between the BimC motor domain and the β-tubulin monomer at 0 mM KCl ion concentration. These salt bridges may play a crucial role in stabilizing BimC binding to tubulins, and subsequently influences the motility of BimC along the microtubule. To get a better visualization of the relative positions of these important salt bridges, these are shown in Figure 6. These high occupancy salt bridges are essential to study the binding mechanism between BimC and the microtubule.

**Figure 6.**
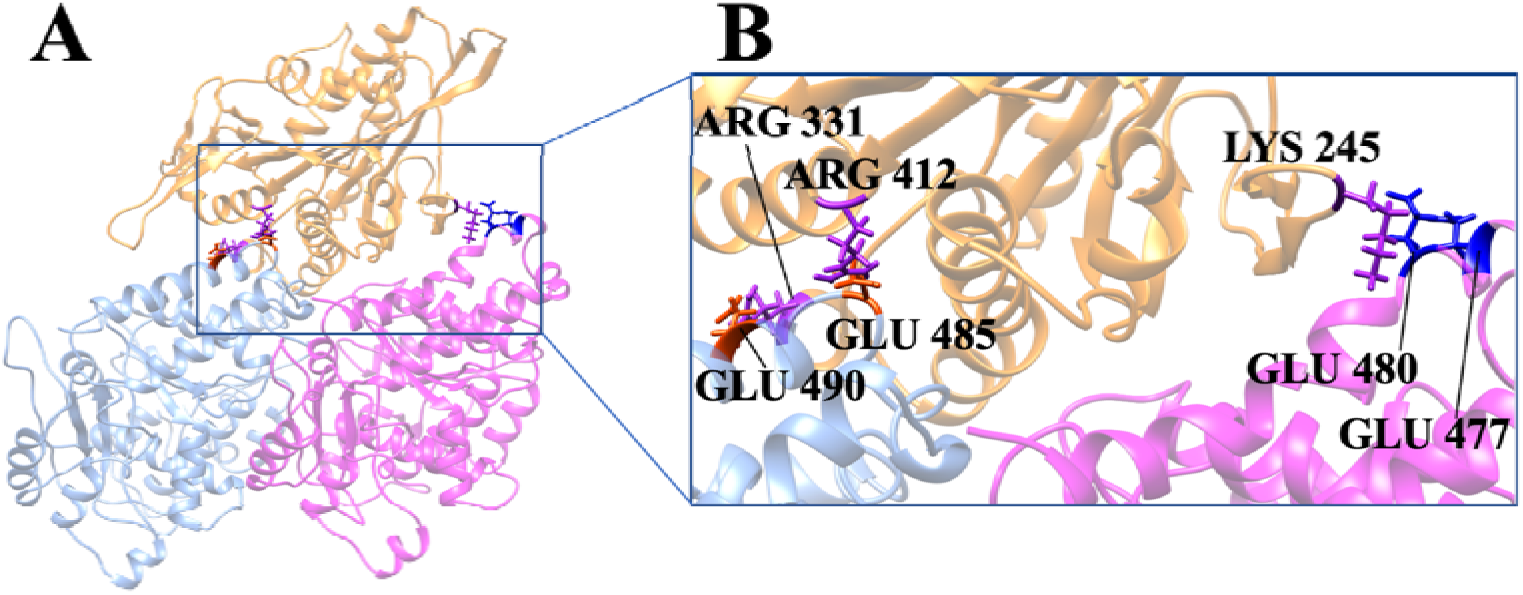
The salt bridges at interfaces between the BimC motor domain and the α/β-tubulin heterodimer. The BimC motor domain, α-tubulin, and β-tubulin are shown in orange, blue, and pink, respectively. Figure B is the close-up view of Figure A.

The positions and occupancies of residues involved in salt bridges on the interfaces between the BimC motor domain and the α/β-tubulin heterodimer at 0 mM KCl concentration are shown in Figure 7. Darker colors represent higher occupancies. The important residues with occupancies above 66% that likely contribute significantly to the binding interactions are marked with their names and occupancies. All the high occupancy residues shown in Figure 7 are on the binding interfaces. Residues ARG331 and ARG412 on the BimC motor domain, and residues GLU490 and GLU485 on α-tubulin, are thought to be essential to stabilize the structure between the BimC motor domain and α-tubulin.

**Figure 7.**
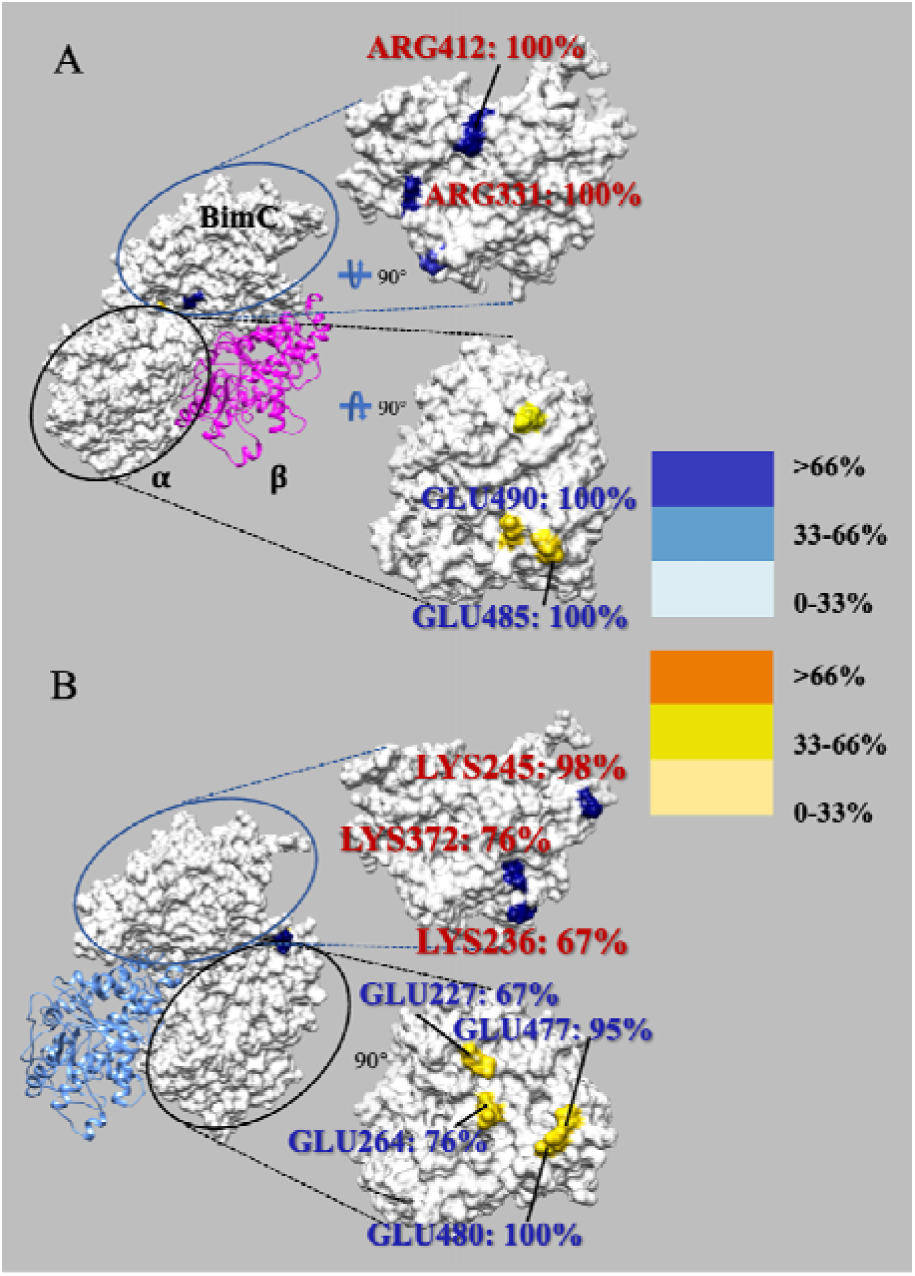
The positions of residues involved in salt bridges between the BimC motor domain and tubulin heterodimer. The important residues with occupancies above 66% are marked with their names and occupancies. (A) shows residues involved in the interaction of the BimC motor domain with α-tubulin. (B) shows residues involved in the interaction of the BimC motor domain with β-tubulin.

Between BimC motor domain and β-tubulin, residues LYS245 on BimC motor domain, and residues GLU480 and GLU477 on β-tubulin, are critical for the complex structure stabilization. These important residues on the BimC motor domain are all positively charged residues, while those on the microtubule are negatively charged, implying that electrostatic interactions are crucial for the binding interaction of BimC motor domains to microtubules.

## Conclusions

The motor domain from *Aspergillus nidulans* BimC is studied in this work. The experimental works were done including generating BimC(Δ1-70) construct and the representative kymograph of BimC(Δ1-70)-GFP at 0/150 mM KCl condition. The result shows BimC walking to the minus end along the microtubule with high velocity at 0mM KCl condition, while there’s no bind between BimC and microtubules at 150 mM KCl concertation. Thus, the experiments indicating that BimC has high binding affinity to microtubules at low ion concertation. It doesn’t bind to microtubules at ion concertation as high as 150 mM KCl. Electrostatic features of BimC and the microtubule at 0 and 150 mM KCl concentrations were investigated using MD simulations and other computational approaches. The motor domain of BimC shows the strongest attractive force at 0 mM KCl to the tubulin heterodimer compared with the complex at 150 mM KCl concentration. The magnitude of electrostatic binding forces shows this pattern clearly with decreased binding forces as the KCl concentration increase. The electrostatic potential also supports this finding. The electrostatic potential shows that the complex structure at 0 mM KCl has the largest surface area of interacting opposite charges compared to the complex at 150 mM KCl. This result corroborates the experimental results, which demonstrates that BimC has higher binding affinity to microtubules at lower ion concentrations and vice versa. Besides, the computational study explained details of this phenomenon.

Furthermore, residues forming salt bridges between the BimC motor domain and tubulin heterodimer at 0 mM KCl concentration were identified by MD simulation. For the two ion concentrations, we find two important salt bridges between the BimC motor domain and α-tubulin, and two between the BimC motor domain and β-tubulin at 0 mM concentration. The two residues (ARG331, ARG412, and LYS245) on BimC which are involved in the important salt bridges could play an important role in nuclear division in Aspergillus. Understanding the nuclear division mechanism may help to design new anticancer drugs. The important residues on the binding interface of BimC are positive, while those residues on the binding interface of the tubulin heterodimer are negative. It indicates that the electrostatic interactions is a key factor of the BimC-microtubule interactions. Deep understanding of the functions of such hot spot residues may help guide kinesin-targeted anticancer drug design [41–46]. Combining computational and experimental results to comprehensively understand the BimC-microtubule interaction is a promising approach to investigating the fundamental mechanisms underlying kinesin dynamics [47].

## Acknowledgment

This research is funded by Grant SC1GM132043 from National Institutes of Health (NIH) and Grant 5U54MD007592 from the National Institutes on Minority Health and Health Disparities (NIMHD), a component of the NIH.

## References

1. Hirokawa, N., et al., Kinesin superfamily motor proteins and intracellular transport. Nature reviews Molecular cell biology, 2009. 10(10): p. 682–696.

2. Kashina, A., G.C. Rogers, and J. Scholey, The bimC family of kinesins: essential bipolar mitotic motors driving centrosome separation. Biochimica et Biophysica Acta (BBA)-Molecular Cell Research, 1997. 1357(3): p. 257–271.

3. Kashina, A., et al., An essential bipolar mitotic motor. Nature, 1996. 384(6606): p. 225.

4. Bishop, J.D., Z. Han, and J.M. Schumacher, The Caenorhabditis elegans Aurora B kinase AIR-2 phosphorylates and is required for the localization of a BimC kinesin to meiotic and mitotic spindles. Molecular biology of the cell, 2005. 16(2): p. 742–756.

5. Enos, A.P. and N.R. Morris, Mutation of a gene that encodes a kinesin-like protein blocks nuclear division in A. nidulans. Cell, 1990. 60(6): p. 1019–1027.

6. Hagan, I. and M. Yanagida, Novel potential mitotic motor protein encoded by the fission yeast cut7+ gene. Nature, 1990. 347(6293): p. 563–566.

7. Le Guellec, R., et al., Cloning by differential screening of a Xenopus cDNA that encodes a kinesin-related protein. Molecular and cellular biology, 1991. 11(6): p. 3395–3398.

8. Blangy, A., et al., Phosphorylation by p34cdc2 regulates spindle association of human Eg5, a kinesin-related motor essential for bipolar spindle formation in vivo. Cell, 1995. 83(7): p. 1159–1169.

9. Guo, W., et al., Using a comprehensive approach to investigate the interaction between Kinesin-5/Eg5 and the microtubule. Computational and structural biotechnology journal, 2022.

10. Ferenz, N.P., A. Gable, and P. Wadsworth. Mitotic functions of kinesin-5. in Seminars in cell & developmental biology. 2010. Elsevier.

11. Drummond, D.R. and I.M. Hagan, Mutations in the bimC box of Cut7 indicate divergence of regulation within the bimC family of kinesin related proteins. Journal of cell science, 1998. 111(7): p. 853–865.

12. Kashlna, A., et al., A bipolar kinesin. Nature, 1996. 379(6562): p. 270–272.

13. Kwok, B.H., J.G. Yang, and T.M. Kapoor, The rate of bipolar spindle assembly depends on the microtubule-gliding velocity of the mitotic kinesin Eg5. Current biology, 2004. 14(19): p. 1783–1788.

14. Walczak, C.E. and T.J. Mitchison, Kinesin-related proteins at mitotic spindle poles: function and regulation. Cell, 1996. 85(7): p. 943–946.

15. Hildebrandt, E.R. and M.A. Hoyt, Mitotic motors in Saccharomyces cerevisiae. Biochimica et Biophysica Acta (BBA)-Molecular Cell Research, 2000. 1496(1): p. 99–116.

16. Kollmar, M. and G. Glöckner, Identification and phylogenetic analysis of Dictyostelium discoideum kinesin proteins. BMC genomics, 2003. 4(1): p. 1–12.

17. Kapitein, L.C., et al., The bipolar mitotic kinesin Eg5 moves on both microtubules that it crosslinks. Nature, 2005. 435(7038): p. 114–118.

18. Sawin, K.E. and T.J. Mitchison, Mutations in the kinesin-like protein Eg5 disrupting localization to the mitotic spindle. Proceedings of the National Academy of Sciences, 1995. 92(10): p. 4289–4293.

19. Dagenbach, E.M. and S.A. Endow, A new kinesin tree. Journal of cell science, 2004. 117(1): p. 3–7.

20. Bevan, D.R., et al., Application of molecular modeling to analysis of inhibition of kinesin motor proteins of the BimC subfamily by monastrol and related compounds. Chemistry & biodiversity, 2005. 2(11): p. 1525–1532.

21. Oguievetskaia, K., et al., Computational fragment-based drug design to explore the hydrophobic sub-pocket of the mitotic kinesin Eg5 allosteric binding site. Journal of computer-aided molecular design, 2009. 23: p. 571–582.

22. Li, L., et al., DelPhi: a comprehensive suite for DelPhi software and associated resources. BMC biophysics, 2012. 5(1): p. 1–11.

23. Li, L., A. Chakravorty, and E. Alexov, DelPhiForce, a tool for electrostatic force calculations: Applications to macromolecular binding. Journal of computational chemistry, 2017. 38(9): p. 584–593.

24. Li, L., et al., DelPhiForce web server: electrostatic forces and energy calculations and visualization. Bioinformatics, 2017. 33(22): p. 3661–3663.

25. Phillips, J.C., et al., Scalable molecular dynamics with NAMD. Journal of computational chemistry, 2005. 26(16): p. 1781–1802.

26. Huszar, D., et al., Kinesin motor proteins as targets for cancer therapy. Cancer and Metastasis Reviews, 2009. 28(1): p. 197–208.

27. Shi, J., J.D. Orth, and T. Mitchison, Cell type variation in responses to antimitotic drugs that target microtubules and kinesin-5. Cancer research, 2008. 68(9): p. 3269–3276.

28. Yu, Y. and Y.M. Feng, The role of kinesin family proteins in tumorigenesis and progression: potential biomarkers and molecular targets for cancer therapy. Cancer, 2010. 116(22): p. 5150–5160.

29. Sanhaji, M., et al., Mitotic centromere-associated kinesin (MCAK): a potential cancer drug target. Oncotarget, 2011. 2(12): p. 935.

30. Hyman, A., Preparation of marked microtubules for the assay of the polarity of microtubule-based motors by fluorescence. Journal of Cell Science, 1991. 1991(Supplement_14): p. 125–127.

31. Roberts, A.J., B.S. Goodman, and S.L. Reck-Peterson, Reconstitution of dynein transport to the microtubule plus end by kinesin. Elife, 2014. 3: p. e02641.

32. Tseng, K.-F., et al., The tail of Kinesin-14a in giardia is a dual regulator of motility. Current Biology, 2020. 30(18): p. 3664–3671. e4.

33. Jumper, J., et al., Highly accurate protein structure prediction with AlphaFold. Nature, 2021. 596(7873): p. 583–589.

34. Bryant, P., G. Pozzati, and A. Elofsson, Improved prediction of protein-protein interactions using AlphaFold2. Nature communications, 2022. 13(1): p. 1–11.

35. Peña, A., et al., Structure of microtubule-trapped human kinesin-5 and its mechanism of inhibition revealed using cryoelectron microscopy. Structure, 2020. 28(4): p. 450–457. e5.

36. Humphrey, W., A. Dalke, and K. Schulten, VMD: visual molecular dynamics. Journal of molecular graphics, 1996. 14(1): p. 33–38.

37. Xian, Y., et al., Structure Manipulation Tool StructureMan: A Structure Manipulation tool to study large scale biomolecular interactions. Frontiers in Molecular Biosciences, 2020. 7: p. 476.

38. Pettersen, E.F., et al., UCSF Chimera—a visualization system for exploratory research and analysis. Journal of computational chemistry, 2004. 25(13): p. 1605–1612.

39. Xian, Y., et al., StructureMan: A structure manipulation tool to study large scale biomolecular interactions. Frontiers in molecular biosciences, 2021. 7: p. 627087.

40. Portnov, I.V. and I.I. Potemkin, Interpolyelectrolyte complex dissociation vs polyelectrolyte desorption from oppositely charged surface upon salt addition. The Journal of Physical Chemistry B, 2020. 124(5): p. 914–920.

41. Sun, S., et al., Electrostatics in Computational Biophysics and Its Implications for Disease Effects. International Journal of Molecular Sciences, 2022. 23(18): p. 10347.

42. Guo, W., et al., A Comprehensive Study on the Electrostatic Properties of Tubulin-Tubulin Complexes in Microtubules. Cells, 2023. 12(2): p. 238.

43. Guo, W., et al., Electrostatic features for nucleocapsid proteins of SARS-CoV and SARS-CoV-2. Mathematical biosciences and engineering: MBE, 2021. 18(3): p. 2372.

44. Warshel, A. and S.T. Russell, Calculations of electrostatic interactions in biological systems and in solutions. Quarterly reviews of biophysics, 1984. 17(3): p. 283–422.

45. Adamczyk, Z., Particle adsorption and deposition: role of electrostatic interactions. Advances in Colloid and Interface Science, 2003. 100: p. 267–347.

46. Sheinerman, F.B. and B. Honig, On the role of electrostatic interactions in the design of protein–protein interfaces. Journal of molecular biology, 2002. 318(1): p. 161–177.

47. Adamczyk, Z. and P. Warszyński, Role of electrostatic interactions in particle adsorption. Advances in Colloid and Interface Science, 1996. 63: p. 41–149.

