## Supplementary material for "How does the ion concentration affect the functions of kinesin BimC": SI

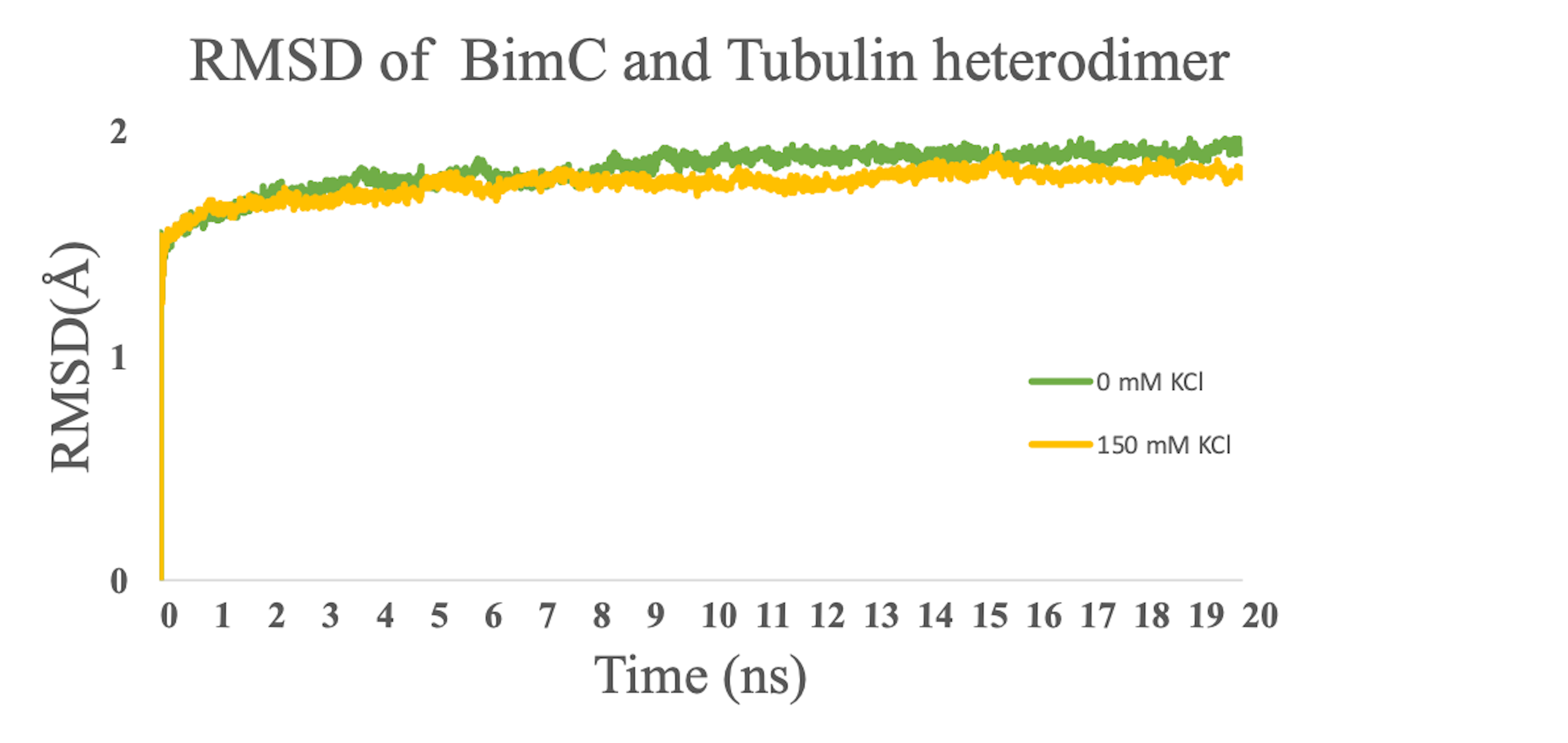

**Figure S1.** The RMSD of BimC-tubulin heterodimer simulations at the two concentrations.

Movies:

Movie 1: The movie corresponds to the kymograph on the left panel in Figure 1a. This movie was acquired with BimC(Δ1-70)-GFP at 0mM KCl condition and single-molecule level motor concentration. Under this condition, BimC(Δ1-70)-GFP exhibited minus-end-directed motility on a single polarity-marked HiLyte 647-microtubule. Top: the microtubule channel, and the arrowhead indicates the microtubule plus end; Middle: the BimC(Δ1-70)-GFP channel; Bottom: the overlay of the microtubule and BimC(Δ1-70)-GFP channels.

Movie 2: The movie corresponds to the kymograph on the right panel in Figure 1a. This movie was acquired with BimC(Δ1-70)-GFP at 150mM KCl condition and very high motor concentration. BimC(Δ1-70)-GFP could not land on the polarity-marked HiLyte 647-microtubule. Top: the microtubule channel; Middle: the BimC(Δ1-70)-GFP channel; Bottom: the overlay of the microtubule and BimC(Δ1-70)-GFP channels.

Movie 3: Simulation of the BimC-tubulin heterodimer (20ns) at 0 mM KCl concentration

Movie 4: Simulation of the BimC-tubulin heterodimer (20ns) at 150 mM KCl concentration
